# *Toxoplasma gondii* and *Trypanosoma lewisi* infection in urban small mammals from Cotonou, Benin, with special emphasis on co-infection patterns

**DOI:** 10.1101/2023.10.15.558972

**Authors:** Jonas R. Etougbétché, Gualbert Houéménou, Antoine A. Missihoun, Philippe Gauthier, Henri-Joël Dossou, Lokman Galal, Ambroise Dalecky, Christophe Diagne, Gauthier Dobigny, Aurélien Mercier

## Abstract

A growing number of studies has highlighted the importance of co-infections in eco-evolutionary processes underlying host-parasite interactions and the resulting epidemiology of zoonotic agents. Small mammals, and particularly rodents, are known to be important reservoirs of many zoonotic pathogens, such as *Toxoplasma gondii* and *Trypanosoma lewisi* that are responsible for toxoplasmosis and atypical trypanosomiasis in human, respectively. Laboratory experiments on rodent models have shown that primary infection with *T. lewisi* increases the host susceptibility to other co-infectious parasites, including *T. gondii*, following an alteration of the immune system. However, data on potential interactions between these parasites in wild small mammals remain scarce. In this study, we estimate the *T. lewisi* prevalence in 553 small mammals from four localities of Cotonou city, Benin. They were then combined with *T. gondii* data previously collected on the same individuals in order to investigate the influence of *T. lewisi* on *T. gondii* infection, and *vice-versa*, using cooccurrence tests and Generalized Linear Mixed Models. Despite quite high overall prevalence (32.5% and 15.2% for *T. gondii* and *T. lewisi*, respectively), we observed a clear and significant segregation between the two parasites. This may be explained by (i) differences in the species-specific susceptibility of small mammal host species to infection by these two parasites, with *R. rattus* and *M. natalensis* being the main reservoirs of *T. lewisi* while *C. olivieri* and *M. m. domesticus* are the main hosts for *T. gondii*; and/or by (ii) a possibly high mortality in co-infected animal in the wild. Although dedicated experimental studies are required to confirm this pattern, as they stand, our data fail to support that infection of small mammals by one of these two parasites favours widespread infection by the second one in nature.

## INTRODUCTION

Within-host interactions between parasites may strongly influence pathobiome dynamics and play a major role in structuring both parasite and host populations [1,2]. Such interactions can have important repercussions on the ecology of zoonotic pathogens, hence on human health, for example by altering the host reservoir susceptibility, modifying the temporal dynamics of infections, increasing transmission risks, or impacting the pathogen virulence [1]. Multi-parasitism is common in all animal organisms, and rodents have been particularly used as model hosts for studies on infection by multiple pathogens [3–6], especially in domestic areas where they are key reservoirs for a wide panel of zoonotic pathogens [3,6,7].

*Toxoplasma gondii* and *Trypanosoma lewisi* are two protozoan parasites of worldwide distribution which are responsible for toxoplasmosis [8] and atypical trypanosomiasis in human [9,10], respectively. Human infection with *T. gondii* usually occurs through consumption of oocyst-contanimated vegetables or undercooked meat, and congenital transmission when primary infection occurs during pregnancy [11]. Toxoplasmosis is usually asymptomatic [12–14] and up to one-third of humans may be infected globally [15–17]. However, clinical forms are sometimes observed, especially in immuno-compromized patients and fetuses [18–21] as well as in immuno-competent individuals from tropical regions infected with *T. gondii* atypical strains that circulate specifically in the environment of these geographical areas [22–24]. *T. lewisi* is transmitted through the feces of infected ectoparasitic fleas that acts as vectors in parasite dissemination among mammals, especially rodents. A few pathogenic and even lethal human infection cases have been described in Asia and Africa; but the global impact of *T. lewisi* on human health may be widely underestimated and remains to be fully documented [9,10,25,26]. In addition to virulence factors of *T. gondii* infecting strain and host-specific genetic factors, co-infections involving *T. gondii* and other parasites are also likely to influence the ecology of toxoplasmosis and host virulence phenotype. For example, the widespread circulation of pathogens such as *T. lewisi* in rodents could increase the infection risk by *T. gondii*, or *vice versa*.

Indeed, experimental studies have shown complex co-infection relationships between *Toxoplasma spp*. and *Trypanosoma spp*. For example *T. lewisi* can have immunosuppressive effects in rats and reduce their immune defenses against co-infectious parasites such as *Toxoplasma gondii* [27–31] whereas in mice, the immune response to *T. gondii* is enhanced following primary infection by *Trypanosoma musculi* [31]. If true in the wild, this would have important consequences for parasite ecology.

Although they display very different transmission modes, previous studies have shown that both parasites can circulate within common environments, especially in tropical areas where they can share the same reservoir hosts, specially rodents [6,32]. Thus, *T. gondii* has been identified in several commensal rodent species [33], including those investigated recently in Cotonou, Benin [34]. Rodents are also the main reservoirs of *T. lewisi* in Africa, especially the invasive genus *Rattus*, which was proposed to play a special role in its ecology and dissemination across the continent [35–38]. Expectedly, it was also detected in many rodents from Cotonou [39]. Both parasites were also found in African shrews of the genus *Crocidura*, in Cotonou city [34,39].

Keeping in mind their usually quite high prevalence in small mammals (e.g., 15.2% for *T. gondii* [34] and 57.2% for *T. lewisi* [39]), the concomitant presence of both parasite in Cotonou city may provide valuable models to investigate further the role of co-infections in eco-evolutionary fate of zoonotic pathosystems in urban reservoir hosts communities.

In this study, we took advantage of an already existing small mammal-borne *Toxoplasma* dataset from Cotonou [34], to assess *Trypanosoma lewisi* presence/absence in the same rodent individuals, and then to investigate specifically the relationships between *Toxoplasma* and

*Trypanosoma* infections in urban wild small mammals taking into account a panel of biological and environmental potentially confounding factors.

## MATERIALS AND METHODS

### Ethical considerations

This study was conducted under the research agreement between the Republic of Benin and the French Institute for Substainable Development (IRD) (renewed on April 6th, 2017) as well as between IRD and Abomey-Calavi University (signed on September 30th, 2010 and renewed on July 3rd, 2019). Field investigations were conducted after a written and/or oral authorization of local authorities (i.e., local heads of urban districts, authorities and staffs of Cotonou Autonomous Seaport) as well as the systematic consent of residents when trapping was conducted inside private settings. Rodents were treated in a human manner according to the American Society of Mammalogy recommendations [40]. None of the species captured in the frame of the current study has IUCN protection status (see CITES list, https://checklist.cites.org/). In line with the Nagoya protocol, authorization for access and equitable sharing of knowledge and data was issued by the competent national authorities of Benin (permit 608/DGEFC/DCPRN/PF-APA/SA).

### Data collection

During 2017 and 2018, a study was conducted on the circulation of *T. gondii* in small mammal from four localities of Cotonou [34]. We here took advantage of these already available small mammal samples to investigate the coinfection patterns between *T. lewisi and T. gondii*. Sampling sites and trapping procedures were previously described in detail [34]. In brief, field campaigns were conducted twice following the same protocols between 2017 and 2018 in three socio-environmentally contrasted districts of the core city, namely Ladji, Agla and Saint-Jean (in October 2017 and June 2018) on the one hand, and in Cotonou seaport (Autonomous Seaport of Cotonou, or ASC) area (in September-November 2017 and March 2018) on the other hand. In each of the three districts, 9-11 households (hereafter designated as “district sites”) were investigated (see details in [39,41]) while nine observatory sites were sampled in ASC (hereafter designated as “ASC sites”; see [42] for their complete description). Small mammals were sexed, age-classed and unambiguously identified at the species level using morphological, DNA sequencing and/or microsatellite genotyping (see details in [34,39,41,42]). We collected different biological samples, including spleen samples that were preserved in 96° ethanol for further molecular assays (see below). The presence of ectoparasitic fleas was checked using fur brushing. We also took advantage of a previous study relying on the same experimental design (i.e., same sampling campaigns, hence same sampling sites) for which the dataset encompassed our own samples in the same districts of Cotonou to obtain socio-environmental data including landcover data, social uses associated with buildings, as well as surface water occurrences. These data comprised 21 GIS-based landscape metrics that were specifically estimated at each district site (see details in [34,43]). Here, we focused on the 553 individuals that had already been investigated for the presence of *T. gondii* using molecular detection (see detailed protocols in [34] and *T. gondii* prevalence data in Table 1 as well as Supplementary Table). Note that these 553 animals are all different from the 369 small mammals used in a previous study on small mammal-borne *T. lewisi* from Cotonou [39] for which no data on *Toxoplasma* were available.

**Table 1:** Prevalence of *T. gondii, T. lewisi* and co-infections by host species and sampled locality. « Tox+ » and « Tryp+ » indicate the number of individuals infected only by *T. gondii* and *T. lewisi*, respectively, while « Tox+ &Tryp+ » correspond to co-infected ones. “Rra”, “Rno”, “Mna”, “Mus”, “Cro, “Cga” and “Pde” stand for *Rattus rattus, R. norvegicus, Mastomys natalensis, Mus musculus domesticus, Crocidura olivieri, Cricetomys gambianus* and *Praomys derooi*, respectively.

| Localities |  | All species | Rra | Rno | Mna | Mus | Cro | Cga | Pde |
| --- | --- | --- | --- | --- | --- | --- | --- | --- | --- |
| Agla | N | **98** | 45 | 11 | 12 | 0 | 30 | 0 | 0 |
|  | Tox+ (%) | **9 (9.2)** | 4 (8.9) | 1 (9.1) | 1 (8.3) | - | 3 (10) | - | - |
|  | Tryp+ (%) | **32 (32.7)** | 23 (51.1) | 0 | 6 (50) | - | 3 (10) | - | - |
|  | Tox+ &Tryp+ (%) | **4 (4.1)** | 2 (4.4) | 0 | 1(8.3) | - | 1 (3.3) | - | - |
| Ladji | N | **91** | 56 | 3 | 3 | 0 | 29 | 0 | 0 |
|  | Tox+ (%) | **12 (13.1)** | 3 (5.4) | 0 | 1 (33.3) | - | 8 (27.6) | - | - |
|  | Tryp+ (%) | **40 (44)** | 32 (57.1) | 2 (66.7) | 2 (66.7) | - | 4 (13.8) | - | - |
|  | Tox+ &Tryp+ (%) | **2 (2.2)** | **0** | 0 | 1 (33.3) | - | 1 (3.4) | - | - |
| St-Jean | N | **73** | 34 | 0 | 7 | 0 | 21 | 7 | 4 |
|  | Tox+ (%) | **14 (19.2)** | 2 (5.9) | - | 1 (14.3) | - | 8 (38.1) | 1 (14.3) | 2 (50) |
|  | Tryp+ (%) | **27 (37)** | 21 (61.8) | - | 3 (42.8) | - | 1 (4.8) | 2 (28.6) | 0 |
|  | Tox+ &Tryp+ (%) | **1 (1.4)** | 1 (2.9) | - | 0 | - | 0 | 0 | 0 |
| ASC | N | **291** | 97 | 52 | 5 | 99 | 38 | 0 | 0 |
|  | Tox+ (%) | **49 (16.8)** | 14 (14.4) | 7 (13.5) | 0 | 20 (20.2) | 8 (21.1) | - | - |
|  | Tryp+ (%) | **82 (28.2)** | 52 (53.6) | 22 (42.3) | 1 (20) | 6 (6.1) | 1 (2.6) | - | - |
|  | Tox+ &Tryp+ (%) | **14 (4.8)** | 7 (7.2) | 5 (9.6) | 0 | 1 (1.0) | 1 (2.6) | - | - |
| All localities | N | **553** | **232** | **66** | **27** | **99** | **118** | **7** | **4** |
|  | Tox+ (%) | **84 (15.2)** | **23 (9.9)** | **8 (12.1)** | **3 (11.1)** | **20 (20.2)** | **27 (22.9)** | **1 (14.3)** | **2 (50)** |
|  | Tryp+ (%) | **181 (32.7)** | **128 (55.2)** | **24 (36.4)** | **12 (44.4)** | **6(6.1)** | **9 (7.6)** | **2 (28.6)** | **0** |
|  | Tox+ &Tryp+ (%) | **21 (3.8)** | **10 (4.3)** | **5 (7.6)** | **2 (7.4)** | **1 (1.0)** | **3 (2.5)** | **0** | **0** |

### Molecular detection of *T. lewisi*

Total genomic DNA was extracted from the spleen using the Qiagen Extraction Kit (DNeasy 96 Blood & Tissue Kit) and according to the manufacturer’s recommendations. DNA elution was performed in 200μL of buffer AE. The presence of *T. lewisi* DNA was checked in independant duplicate through a qPCR protocol previously described [35,38]. The latter procedure targets a 131 bp-long fragment of the *Trypanosoma* 18S rDNA gene fragment, using primers Trypano 1 (5’-AGGAATGAAGGAGGGTAGTTCG-3’) and Trypano 2 (5’-CACACTTTGGTTCTTGATTGAGG-3’) as well as two hybridization probes (Trypano 3: [LC640] AGAATTTCACCTCTGACGCCCCAGT [Phos], and Trypano 4: GCTGTAGTTCGTCTTGGTGCGGTCT [Flc]). Amplification conditions included a denaturation phase at 95°C for 10 min, followed by 60 cycles (95°C for 10s, 56°C for 15s then 72°C for 20s) and a cooling phase at 40°C for 10s performed on a LightCycler 480 (Roche) using a 384-well microtiter plate and 3μL of DNA template in a final volume of 10μL for each reaction. Genomic DNA extracts from *T. lewisi* and *T. brucei* cell cultures were used as positive controls, while sterile water served as a negative control. The sigmoidal shape of each amplification curve was checked by-eye in order to discard non-sigmoidal signals that may represent false positive results. All individuals that provided at least one positive signal (out of the two duplicate qPCR experiments) were considered *Trypanosoma*-positive. The qPCR results were expressed as cycle threshold values. Note that, *Trypanosoma*-positive samples with sufficient DNA (Ct ≤ 30) were genotyped relying on 9 *T. lewisi*-specific microsatellite markers recently developed by [44] for unambiguous *T. lewisi* molecular identification (data not shown).

### Data analysis

Chi-square tests were used to compare parasite prevalence between host species and/or between trapping localities. We carried out these tests for *T. lewisi* prevalence (between host species and between localities) on the one hand, and for *T. gondii – T. lewisi* co-infection (between and within host species) on the other hand. We then performed two complementary sets of analyses to explore the possible interactions between both parasites in relation to host-intrinsic and host-extrinsic factors: co-occurrence analyses and Generalized Linear Mixed Models (GLMMs).

### ‘Co-occurrence analyses’

Co-occurrence analyses are used to test whether two entities are found statistically aggregated or statistically segregated more often than expected under random association [45,46]. Here, such deterministic *vs*. random associations of the two parasite species were tested depending on the host species or on the locality, as well as on the whole dataset combining all host species and localities. To do so, data were organized in several matrices following the different small mammal species and different sampled localities: each column corresponded to a host individual while each row indicated the absence (0) or the presence (1) of a given parasite species (i.e., one row for *T. lewisi*, another one for *Toxoplasma*). Only matrices with at least 10 host individuals were considered in this analysis. Observed data were compared to expected results under the null hypothesis of random assembly with a 95% confidence limit [45,47] in PAIRS v.1.0 [48] and using the standardized Z-score (ZCS) [49] as a quantitative index of co-occurrence. Significant negative and positive ZCS indicated aggregation and segregation, respectively [45]. Statistical significance was assessed by comparing the observed ZCS to values obtained from 10,000 iterations using a statistically recommended null model using the fixed row and equiprobable column constraints algorithm [45].

### ‘Generalized Linear Mixed Models (GLMM)’

Generalized Linear Mixed predictive Models (GLMMs) were tested not only on the whole small mammal community, but also for each species with at least 50 individuals sampled, in order to explore the relationships between the prevalence of *T. gondii* and that of *T. lewisi* in small mammals across Cotonou city, taking geographic/environmental parameters into account. These analyses were performed separately for the three urban districts (herefater designed as to ‘district-based models’) on the one hand, and for the ASC (‘ASC-based models’) on the other hand, since (i) these two areas display very distinct socio-economic, historical and environmental characteristics, (ii) no landscape/GIS data was available for the ASC, and (iii) the trapping campaigns were not carried out exactly at the same period in the seaport and the core city (see above). For each dataset, three models were tested with (1) the prevalence of *T. gondii*, (2) the prevalence of *T. lewisi* and (3) the prevalence of co-infections as binary response variables, respectively.

In each model, we considered the individual characteristics of the host (sex, age and presence/absence of fleas), the period of capture (i.e., trapping session) and socio-environmental proxies (i.e., trapping sites coordinates along the first four Principal Components retrieved from the set of 21 GIS-based landscape metrics and available only for district-based models) as explanatory variables. For the first two models, when the prevalence of *T. lewisi* was used as response variable, that of *T. gondii* was added to explanatory variables, and *vice versa*. Districts (in district-based models) and ASC sites (in ASC-based models) were considered as random variables in order to account for possible spatial variation or autocorrelation.

Models with all possible combinations of the terms included in the starting model were generated, and the most parsimonious model (i.e., the one explaining the highest variance level with the fewest explanatory variables) was chosen among those selected within two AIC units of the best model retrieved [50]. The significance of explanatory variables and their interactions was determined by deletion testing and log-likelihood ratio tests and, when needed, by pairwise Wilcoxon rank sum tests with 95% family-wise confidence level. The final model was validated by the over-dispersion test, the graphical checking of normality, independence as well as variance homogeneity of residuals. These analyses were performed in R [51] using dedicated packages, namely *lme4* for GLMMs [52] and *MuMIN* for model selection [53].

## RESULTS

### Sampling

The two parasites *T. gondii* and *T. lewisi* could be concomittantly investigated in 553 individuals: 232 black rats (*Rattus rattus*), 118 African giant shrews (*Crocidura olivieri*), 99 house mice (*Mus musculus domesticus*), 66 brown rats (*Rattus norvegicus*), 27 multi-mammate rats (*Mastomys natalensis*), 7 Gambian pouched rats (*Cricetomys gambianus*) and 4 Deroo’s mice (*Praomys derooi*). Among them, there were 406 adults and 115 juveniles (32 individuals displayed ambiguous patterns and their age could not be assessed with confidence), 245 males and 308 females. Eighty out of 553 (i.e 14.5%) animals carried at least one flea with the highest pulicidal prevalence observed in *Rattus norvegicus* (50%, i.e 33 flea-carrying individuals out of 66). Captures per locality were distributed as follows: 98 small mammals in Agla (45 *R. rattus*, 30 *C. olivieri*, 12 *M. natalensis*, 11 *R. norvegicus*), 91 in Ladji (56 *R. rattus*, 29 *C. olivieri*, 3 *M. natalensis*, 3 *R. norvegicus*), 73 in Saint-Jean (34 *R. rattus*, 21 *C. olivieri*, 7 *M. natalensis*, 7 *C. gambianus*, 4 *P. derroi*) and 291 in ASC (99 *M*.*m. domesticus*, 97 *R. rattus*, 52 *R. norvegicus*, 38 *C. olivieri*, 5 *M. natalensis*) (see Figure 1 and Table 1 for details).

**Figure 1:**
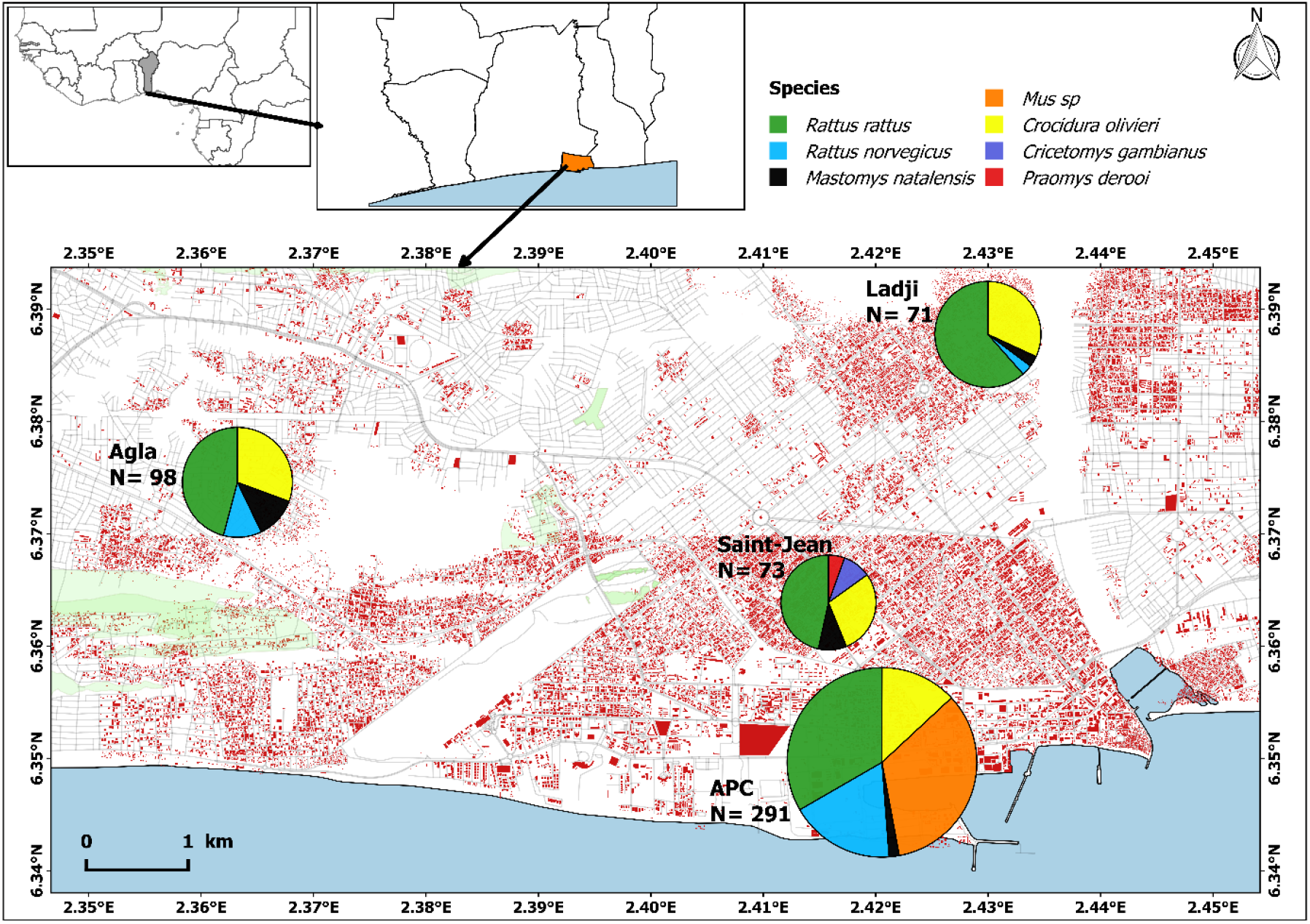
Species-specific distributions and relative abundances of the samples available for the present study (i.e., that were investigated for both parasites). Modified from Etougbétché *et al*. [34].

### Small mammal-borne *T. lewisi* prevalence

Out of the 553 individuals screened, a total of 181 individuals were found *T. lewisi*-positive, thus representing an overall molecular prevalence of 32.7% (see details in Table 1). The highest prevalence was found in *R. rattus, M. natalensis* and *R. norvegicus* with 55.2% (128/232), 44.4% (12/27) and 36.4% (24/66), respectively, with much lower prevalence in *M. m. domesticus* (6/99; 6.1%) and *Crocidura olivieri* (9/118; 7.6%), respectively. A significant difference in *T. lewisi* infection was observed between host species (χ2 = 120.33, df = 4, *p* < 10^-3^) with black rats being more infected than other species (*p* < 0.01), except *M. natalensis* (χ2 = 0.73, df = 1, *p* = 0.42). *C. gambianus* and *P. derooi* showed only one and no individual infected, respectively; however, they were very poorly represented in our dataset (n=7 and n=4) and were thus removed from subsequent analyses. Prevalences were significantly different between localities (χ2 = 8.5, df = 3, *p* = 0.036), with small mammals from Ladji being the most infected ones (40/91; 44%), followed by those from St-Jean (27/73; 37%), Agla (32/98; 32.7%) and ASC (82/291; 28.2%).

### Host species-specific prevalence of *T. gondii – T. lewisi* co-infection

Only 21 out of 553 (3.8%) of the studied individuals were found infected with both parasites. Among these co-infected individuals, no significant differences (χ^2^ = 1.66, df = 1, *p* = 0.19 were found between *R. rattus* (10/21; 47.7%) and *R. norvegicus* (5/21; 23.8%), but black rats were significantly more co-infected than the other species (χ^2^, all *p* < 0.04) while no difference was found between *R. norvegicus* and the other species where rather low prevalence were found: *C. olivieri* (14.28%, i.e 3/21) and *M. natalensis* (9.52%, i.e 2/21). Only one (1/99) *M. m. domesticus* was found infected with both parasites. Comparison of species-specific co-infection prevalence between localities have shown no significative difference in *R. rattus* (Fisher’s Exact, *p* = 0.17) and in *C. olivieri* (Fisher’s Exact, *p* = 1). All co-infected *R. norvegicus* were found in ASC, thus precluding any inter-locality investigation for this particular species.

### Investigation of *T. gondii* - *T. lewisi* co-infection patterns

#### ‘Co-occurrence analysis’

Most of the tests for co-occurrence of *T. gondii* and *T. lewisi* showed significant segregation between the two parasites, at both the host species and locality levels (all standardized Z-scores > 0 and all *p*-values ≤ 0.002, except for *M. natalensis* that showed only marginally non-significant probability; Tab. 2). Considering the whole small mammal dataset at the scale of Cotonou city the segregation pattern was also highly significant (Tab. 2).

#### ‘GLMM analysis’

Although several predictive models were tested (see Table 3), in no instance did we find that the infection by one of the two parasites could be explained by the infection by the other one. This was true whatever the design of the model, the host species and the considered area.

**Table 2:** Co-occurrence of *T. gondii* and *T. lewisi* in small mammal species and sampled localities. N: Number of individuals infected by at least one parasite; Try+ and Tox+: number of individuals infected only by *T. lewisi* and *T. gondii*, respectively; Tox+&Tryp+: number of individuals in which both parasites were detected. ZCS : standardized Z-score.

|  |  | N | Try+ | Tox+ | Tox+&Tryp+ | ZCS | *p* |
| --- | --- | --- | --- | --- | --- | --- | --- |
| Species | *R. rattus* | 140 | 127 | 23 | 10 | 9.39 | <0.001 |
|  | *R. norvegicus* | 27 | 24 | 8 | 5 | 3.05 | 0.002 |
|  | *M. natalensis* | 13 | 12 | 3 | 2 | 1.83 | 0.067 |
|  | *Mus m. domesticus* | 24 | 20 | 5 | 1 | 4.73 | <0.001 |
|  | *C. olivieri* | 33 | 27 | 9 | 3 | 5.12 | <0.001 |
| Localities | Agla | 36 | 31 | 9 | 4 | 4.62 | <0.001 |
|  | Ladji | 50 | 40 | 12 | 2 | 7.53 | <0.001 |
|  | St-Jean | 40 | 27 | 14 | 1 | 8.13 | <0.001 |
|  | ASC | 116 | 81 | 49 | 14 | 10.96 | <0.001 |
| Total |  | 242 | 179 | 84 | 21 | 16.45 | <0.001 |

**Table 3:**
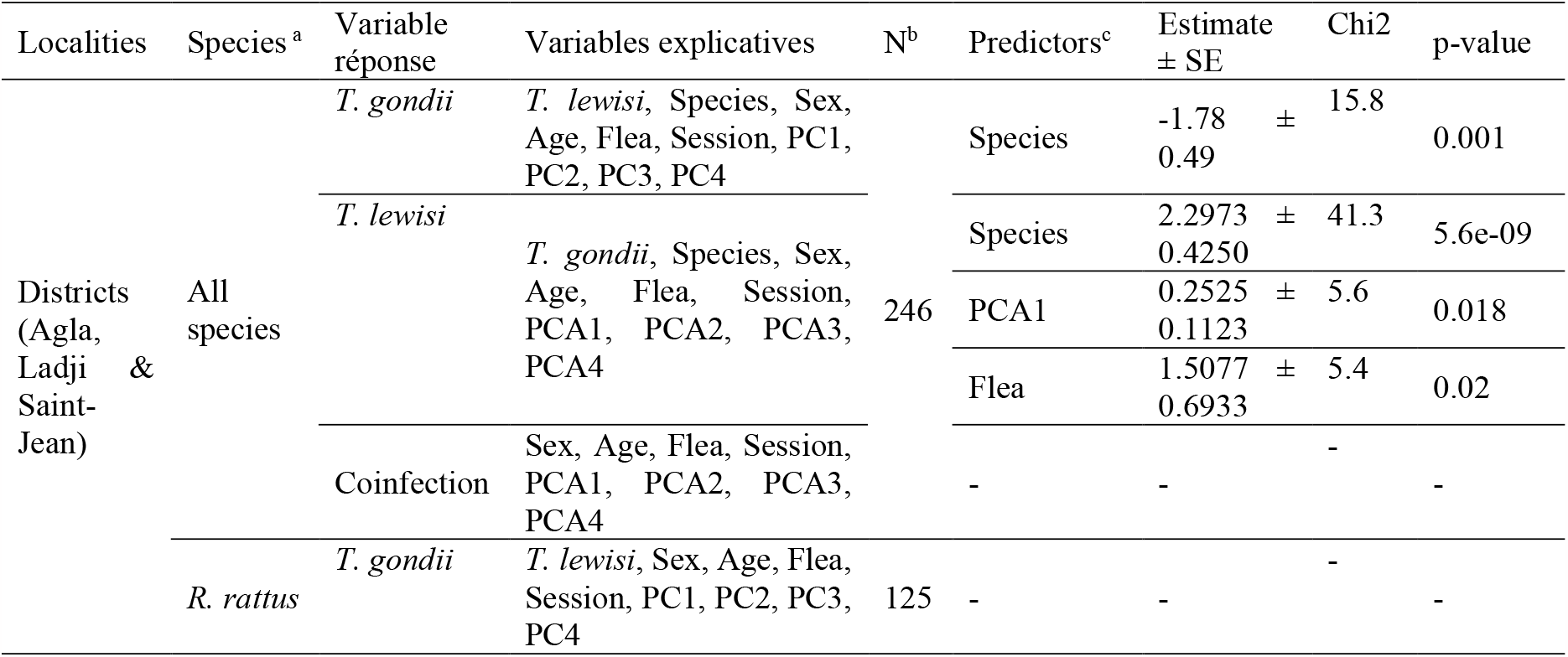
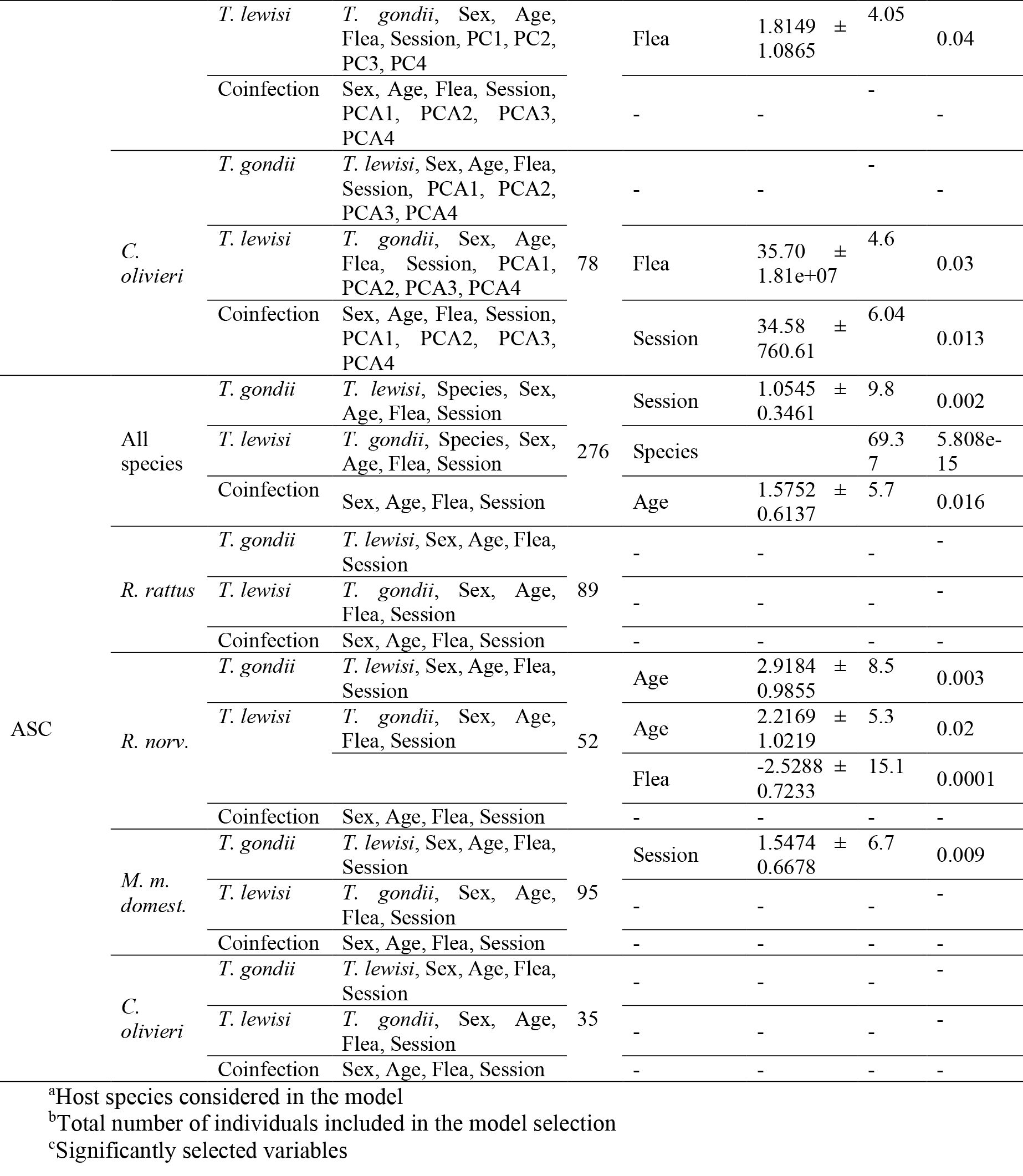
Results of the best-fitted Generalized Linear Mixed Models explaining *T. gondii* and *T. lewisi* mono-infections as well as co-infections in the different small mammal populations/species sampled. Flea: presence of fleas on the individual. Session: period of capture. PC 1 to 4: sites coordinates on landscape PCA axes 1 to 4 (see text and Dossou et al.[43] for details).

##### - District-based analysis

Considering all host species across the three urban districts, none of the two parasite species could explain the presence of the other. Examining co-infected species-specific patterns, only co-infection in *C. oliveri* seemed to be related to the trapping session (χ2 = 6.04; *p =* 0.014): they were significantly more commonly co-infected in October 2017 than in June 2018. Note that infection with any of the two parasites was significantly related to host species. Black rats were significantly less infected by *T. gondii* than other species (χ2 = 15.79; *p =* 0.001; Wilcoxon test, *p* < 0.001) while *T. lewisi* infection varied between host species (χ2 = 41.29; *p* < 0.001): pairwise comparison of *T. lewisi* infection showed that *R. rattus* was significantly more infected than *C. olivieri* and *R. norvegicus*, whereas *M. natalensis* was more infected than *C. olivieri* (Wilcoxon test, all *p*-values < 0.01). No difference in infection between *R. rattus* and *M. natalensis* was observed (*p* = 0.75). In addition to host species, *T. lewisi* infection was also positively associated with the presence of ectoparasitic fleas (χ2 = 5.42; *p* = 0.02) as well as partly with the landscape structure, namely PC1 which contrast dumpsites to houses (χ2 = 5.6; *p* < 0.017). Focusing on the two best represented host species in our dataset, namely *Rattus rattus* and *Crocidura olivieri*, we confirmed that only *T. lewisi* infection was significantly associated with the presence of fleas (χ2 = 4.05; *p* = 0.04 and χ2 = 4.64; *p* = 0.03 respectively).

##### - ASC-based analyses

As for the urban districts, no statistically significant relationship between the two parasites was observed in ASC, whatever the host species considered. However, when considering all host species, the most parcimonious model best explaining co-infection included the age stage (χ2 = 5.7; *p* = 0.016), with juveniles being more co-infected than adults. Furthermore, when all host species were considered, *T. lewisi* infection was also found significantly related to the host species in ASC (χ2 = 69.37; *p* < 0. 001). The genus *Rattus* was once again found as the most infected one (*R. rattus* vs. *C. olivieri* /*M. m. domesticus*, Wilcoxon tests, both *p* < 0.001; *R. norvegicus* vs. *C. olivieri*/*M. m. domesticus*, Wilcoxon tests, both *p* < 0.001).

## DISCUSSION

Our study confirms the role of commensal small mammals in the large-scale circulation of two environnemental parasites within Cotonou City, namely *Trypanosoma lewisi* and *Toxoplasma gondii* with overall molecular prevalences of 32.7% and 15.2%, respectively. Both parasites were observed in all investigated localities, although the level of their respective prevalence is variable from one to another. The implications of intrinsic and extrinsic factors considered in this study in *T. gondii* infections has been extensively discussed in our previous study [34]. For this reason, beyond the socio-environmental patterns that explain concomitant presence of both parasites in some rodent and shrew individuals which is the main focus of the present study, we first discuss briefly some aspects of *T. lewisi* infections as well.

Our results are quite congruent with previous studies that already showed that *Trypanosoma lewisi* was widespread among domestic and peri-domestic small mammals, with the black rat being the most widespread and important reservoir species in this part of West Africa (e.g., [35,38], including in Cotonou city [39]. As such, the overall qPCR-based prevalence observed within Cotonou by Dobigny and colleagues [39] and the present study were 57.2% (66.9% in black rats) and 32.7% (55.2% in black rats), respectively. However, contrary to Dobigny and colleagues study [39], we here found that *M. natalensis*-specific prevalence (44.4%) was not statistically different from that observed in black rats, even if ranges observed in the two studies remain quite similar (33.9% in *M. natalensis* in [39]). This observation seems to show that, in addition to the invasive genus *Rattus* usually considered as the main reservoir of *T. lewisi*, the native *M. natalensis* may also play a major eco-epidemiological role in the maintenance and circulation of *T. lewisi* in urban environments. Species-specific prevalences observed in Cotonou appear higher than those retrieved from other African contexts using the same molecular detection technic: e.g., 14.4% in Uganda (29.5% in black rats; [37]), 14.6% in Niger and Nigeria (25.2% in black rats; [38]) or 15.5% in Senegal (27.8% of black rats; [54]). In our study, we detected no significant association between landscape and small mammal-borne *T. lewisi*, thus suggesting that this particular parasite may be widely distributed in most of the city. Considering the district-based analysis, the GLMM performed on all species as well as those performed on the two most represented species, showed that *T. lewisi* infection was related not only to the host species (with rats as the main reservoirs) but also to the presence of fleas on the animals. The latter observation is not surprising since fleas are the main transmission vectors of *T. lewisi* in rodents in general, and particulary the genus *Rattus* [36,55,56].

Although both parasites were observed in Cotonou small mammal community (15.2% for *T. gondii* and 38.7% for *T. lewisi*), there was no sign of association, i.e. favoured co-infection. GLMM-based analyses showed no statistically significant relationship between both parasites species regardless of the strategies used. This supports the absence of co-infections that would be favored. By the contrary, a clear trend towards segregation was even observed by our co-occurrence analyses regardless of the host species (i.e., all or individual ones) and the locality considered. Our results are in strong contradiction to what was expected from experimental data available in literature, particulary in rats, which suggest that primary infection by *T. lewisi* would compromise host immunity and either favour infection by *T. gondii*.

The segregation between the two parasites that we observed on the field could be explained by the species-specific composition of small mammals in the study sites, together with differences in the host-species specific susceptibility to these two parasites. Indeed, contrast, black rats appear less susceptible to *T. gondii* infection than shrews and house mice, in which significantly higher prevalence levels were observed in south Benin [34]. Unfortunately, robust data on virulence of these two parasites in African rodents are not available, thus precluding any definitive conclusion.

The rarity of *T. gondii* - *T. lewisi* co-infected animals in our dataset (3.8%: 21 out of 553) could also be explained by a differential mortality that would limit our ability to detect the simultaneous presence of both parasites in the field. Indeed, if infection with *T. lewisi* leads to a severe alteration of the immune system, a second infection with *T. gondii* could be lethal, thus reducing the lifespan of co-infected animals. If this was to be true co-infection rates observed in natural small mammal populations would be very reduced. At the same time, Gao and colleagues, [31] found that laboratory Norway rats infected with *T. lewisi* were more susceptible to *T. gondii* infections. Although this does not necessarily remain true for wild rodents, *a fortiori* for wild black rats, a *T. lewisi*-induced increased susceptibility to *T. gondii* would be expected to an increase in the proportions of co-infected animals [27–31]. Yet, this was not observed, thus rather suggesting very high mortality in co-infected animals, or no co-infection. On the sole basis of our data, and in absence of experimental data on coinfection-associated mortality, it appears difficult to decide between the two explanations.

Indeed, several aspects may have weakened our study and the interpretation of its results. First, in murine models, experimental infestations have demonstrated different levels of susceptibility/resistance to *T. gondii* between species or even lineages of the same species, which may depend on the host genetic background and/or the particular strain of *T. gondii* used for inoculations [60,61]. Unfortunately, no such data are available for wild rodents and circulating *T. gondii* strains in Benin. Second, under natural conditions, knowing in which rank parasites infected one given host is not trivial, if not impossible. Yet, it is very likely that a *T. gondii* infection prior to that of *T. lewisi* will not induce the same immune response, hence will not have the same physiological consequences than a *T. lewisi* infection followed by a *T. gondii* infection. More generally, during their lifespan, wild rodents are very likely to be infected by several pathogens, some of them potentially strongly impacting their immune system and general condition, hence the fate of subsequent encounters with other pathogens. This of course may strongly obscure the study of co-infection patterns. For example, several studies have suggested that infection with *T. lewisi* may induce an immunosuppressive effect on its hosts, leading to increased susceptibility to infection by *T. gondii*, but also by other pathogens such as bacteria, viruses, helminths and protozoa [1,2,59–64]. This suggests that study of co-infections in wild hosts should include the investigation of large panels of pathogens and parasites (e.g., through digestive tract analyses, highthrouput DNA sequencing approaches, etc). In turn, this also implies to rely on very large amount of hosts in order to reach sufficient statistical power. Alternatively, experimental co-infections on wild or wild-derived rodents may allow one to investigate simplified co-infection processes and eco-evolutionary consequences in better controlled/documented frameworks.

In conclusion, our study provides new insights into the interactions *in natura* between two urban small mammal-borne parasites with zoonotic potential in Africa, particularly in Benin. We confirmed the extensive circulation of *T. lewisi* among domestic small mammals within Cotonou city, especially in the abundant and widespread *R. rattus*, thus confirming potential spillover risk to city dwellers [39]. We also observed a statistically significant segregation between *T. gondii* and *T. lewisi* in their hosts and failed to detect any evidence that the infection by one of these two parasites may favour the infection by the other in the wild, as could have been expected from experimental studies previously conducted on laboratory rats. An unfrequently co-infection in Cotonou city was also observed, potentially due to differences in the susceptibility of host species to infection by these two parasites, and/or by a high mortality of co-infected individuals in the wild which would preclude their detection on the field. Experimental studies on wild rodent models are required to document further these two hypotheses. However, our results strongly suggest that, whatever the underlying process, coinfection by one of the two parasite is not a major driver of widescale and persistent infection by the second one in rodents.

## Acknowledgements

This study is part of a long-term partnership between Cotonou Autonomous Seaport, the Polytechnic School of Abomey-Calavi, the French Institute of Research for Sustainable Development, and the Tropical Neurology Institute (Inserm U1094, IRD U270 EpiMaCT, University of Limoges). We are grateful to Ladji, Agla and Saint-Jean authorities as well as inhabitants who kindly authorized us to access their households for trapping and interview purposes. We thank the Autonomous Port of Cotonou authorities and staff who facilitated our access to their infrastructures. We also thank the CBGP Small Mammal Collection (Centre de Biologie pour la Gestion des Populations, 2018, “CBGP – Small mammal Collection”, https://doi.org/10.15454/WWNUPO) for the conservation of samples from Benin.

## Authors contributions

**Design of the study:** A. Mercier and G. Dobigny; **Sampling**: J. R. Etougbétché, H.-J. Dossou, and G. Dobigny; **Molecular biology work**: J. R. Etougbétché and P. Gauthier; **Data analysis and processing**: J. R. Etougbétché, C. Diagne, A. Dalecky; **Paper writing & review**: J. Etougbétché, Lokman Galal, C. Diagne, G. Dobigny, A. Mercier. **Supervision**: A. Mercier, G. Dobigny, G. Houéménou, A. A. Missihoun and I. Youssao Abdou Karim.

---

**Supplementary Table:**
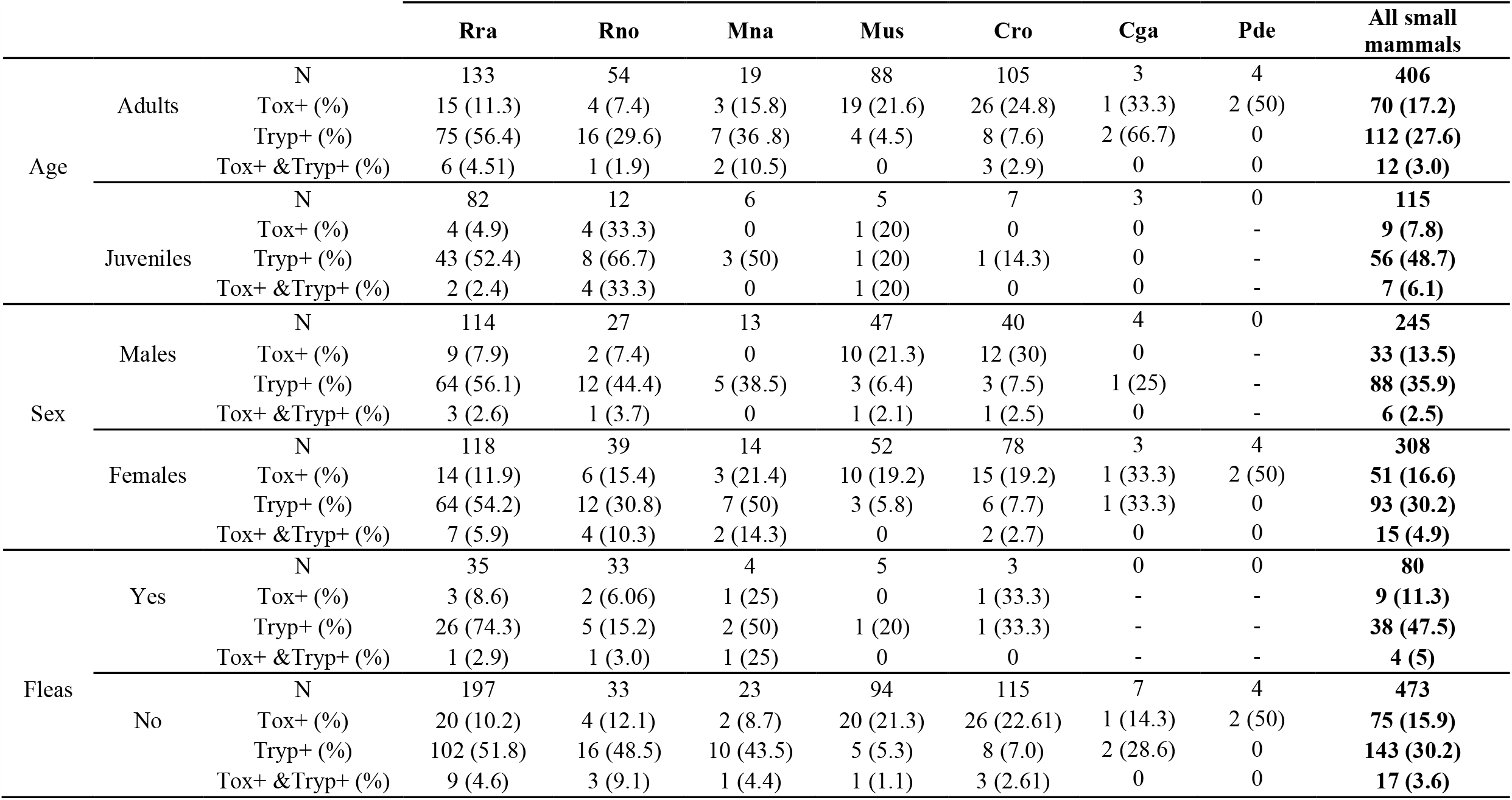
Prevalence of *T. gondii, T. lewisi* and Co-infection by sex, age, session and fleas porting. « Tox+ » and « Tryp+ » indicate the number of *T. gondii-* and *T. lewisi* infected individuals, respectively, while « Tox+ &Tryp+ » correspond to co-infected ones. “Rra”, “Rno”, “Mna”, “Mus”, “Cro, “Cga” and “Pde” stand for *Rattus rattus, Rattus norvegicus, Mastomys natalensis, Mus musculus domesticus, Crocidura olivieri, Cricetomys gambianus* and *Praomys derooi*, respectively.

|  |  |  | **Rra** | **Rno** | **Mna** | **Mus** | **Cro** | **Cga** | **Pde** | **All small mammals** |
| --- | --- | --- | --- | --- | --- | --- | --- | --- | --- | --- |
| Age | Adults | N | 133 | 54 | 19 | 88 | 105 | 3 | 4 | **406** |
|  |  | Tox+ (%) | 15 (11.3) | 4 (7.4) | 3 (15.8) | 19 (21.6) | 26 (24.8) | 1 (33.3) | 2 (50) | **70 (17.2)** |
|  |  | Tryp+ (%) | 75 (56.4) | 16 (29.6) | 7 (36.8) | 4 (4.5) | 8 (7.6) | 2 (66.7) | 0 | **112 (27.6)** |
|  |  | Tox+ &Tryp+ (%) | 6 (4.51) | 1 (1.9) | 2 (10.5) | 0 | 3 (2.9) | 0 | 0 | **12 (3.0)** |
|  | Juveniles | N | 82 | 12 | 6 | 5 | 7 | 3 | 0 | **115** |
|  |  | Tox+ (%) | 4 (4.9) | 4 (33.3) | 0 | 1 (20) | 0 | 0 | - | **9 (7.8)** |
|  |  | Tryp+ (%) | 43 (52.4) | 8 (66.7) | 3 (50) | 1 (20) | 1 (14.3) | 0 | - | **56 (48.7)** |
|  |  | Tox+ &Tryp+ (%) | 2 (2.4) | 4 (33.3) | 0 | 1 (20) | 0 | 0 | - | **7 (6.1)** |
| Sex | Males | N | 114 | 27 | 13 | 47 | 40 | 4 | 0 | **245** |
|  |  | Tox+ (%) | 9 (7.9) | 2 (7.4) | 0 | 10 (21.3) | 12 (30) | 0 | - | **33 (13.5)** |
|  |  | Tryp+ (%) | 64 (56.1) | 12 (44.4) | 5 (38.5) | 3 (6.4) | 3 (7.5) | 1 (25) | - | **88 (35.9)** |
|  |  | Tox+ &Tryp+ (%) | 3 (2.6) | 1 (3.7) | 0 | 1 (2.1) | 1 (2.5) | 0 | - | **6 (2.5)** |
|  | Females | N | 118 | 39 | 14 | 52 | 78 | 3 | 4 | **308** |
|  |  | Tox+ (%) | 14 (11.9) | 6 (15.4) | 3 (21.4) | 10 (19.2) | 15 (19.2) | 1 (33.3) | 2 (50) | **51 (16.6)** |
|  |  | Tryp+ (%) | 64 (54.2) | 12 (30.8) | 7 (50) | 3 (5.8) | 6 (7.7) | 1 (33.3) | 0 | **93 (30.2)** |
|  |  | Tox+ &Tryp+ (%) | 7 (5.9) | 4 (10.3) | 2 (14.3) | 0 | 2 (2.7) | 0 | 0 | **15 (4.9)** |
| Fleas | Yes | N | 35 | 33 | 4 | 5 | 3 | 0 | 0 | **80** |
|  |  | Tox+ (%) | 3 (8.6) | 2 (6.06) | 1 (25) | 0 | 1 (33.3) | - | - | **9 (11.3)** |
|  |  | Tryp+ (%) | 26 (74.3) | 5 (15.2) | 2 (50) | 1 (20) | 1 (33.3) | - | - | **38 (47.5)** |
|  |  | Tox+ &Tryp+ (%) | 1 (2.9) | 1 (3.0) | 1 (25) | 0 | 0 | - | - | **4 (5)** |
|  | No | N | 197 | 33 | 23 | 94 | 115 | 7 | 4 | **473** |
|  |  | Tox+ (%) | 20 (10.2) | 4 (12.1) | 2 (8.7) | 20 (21.3) | 26 (22.61) | 1 (14.3) | 2 (50) | **75 (15.9)** |
|  |  | Tryp+ (%) | 102 (51.8) | 16 (48.5) | 10 (43.5) | 5 (5.3) | 8 (7.0) | 2 (28.6) | 0 | **143 (30.2)** |
|  |  | Tox+ &Tryp+ (%) | 9 (4.6) | 3 (9.1) | 1 (4.4) | 1 (1.1) | 3 (2.61) | 0 | 0 | **17 (3.6)** |

## REFERENCES

1. Vaumourin E, Vourc’h G, Gasqui P, Vayssier-Taussat M. The importance of multiparasitism: examining the consequences of co-infections for human and animal health. Parasit Vectors. 2015;8:545.

2. Venter F, Matthews KR, Silvester E. Parasite co-infection: an ecological, molecular and experimental perspective. Proc R Soc B. 2022;289:20212155.

3. Anh LTL, Balakirev AE, Chau NV. Investigation of multiple infections with zoonotic pathogens of rodents in northern Vietnam. J Vector Borne Dis. 2021;58:47–53.

4. Costa F, Porter FH, Rodrigues G, Farias H, de Faria MT, Wunder EA, et al. Infections by Leptospira interrogans, Seoul virus, and Bartonella spp. among Norway rats (Rattus norvegicus) from the urban slum environment in Brazil. Vector-Borne Zoonotic Dis. 2014;14:33–40.

5. Diagne C, Galan M, Tamisier L, d’Ambrosio J, Dalecky A, Bâ K, et al. Biological invasions in rodent communities: from ecological interactions to zoonotic bacterial infection issues. bioRxiv. 2017;108423.

6. Schmidt S, Essbauer SS, Mayer-Scholl A, Poppert S, Schmidt-Chanasit J, Klempa B, et al. Multiple infections of rodents with zoonotic pathogens in Austria. Vector Borne Zoonotic Dis Larchmt N. 2014;14:467–75.

7. Han BA, Kramer AM, Drake JM. Global Patterns of Zoonotic Disease in Mammals. Trends Parasitol. 2016;32:565–77.

8. de Barros RAM, Torrecilhas AC, Marciano MAM, Mazuz ML, Pereira-Chioccola VL, Fux B. Toxoplasmosis in Human and Animals Around the World. Diagnosis and Perspectives in the One Health Approach. Acta Trop. 2022;231:106432.

9. Kumar R, Gupta S, Bhutia WD, Vaid RK, Kumar S. Atypical human trypanosomosis: Potentially emerging disease with lack of understanding. Zoonoses Public Health. 2022;69:259–76.

10. Truc P, Büscher P, Cuny G, Gonzatti MI, Jannin J, Joshi P, et al. Atypical human infections by animal trypanosomes. PLoS Negl Trop Dis. 2013;7:e2256.

11. Dubey JP. Toxoplasmosis of animals and humans. CRC press; 2016.

12. Attias M, Teixeira DE, Benchimol M, Vommaro RC, Crepaldi PH, De Souza W. The life-cycle of Toxoplasma gondii reviewed using animations. Parasit Vectors. 2020;13:1–13.

13. Halonen SK, Weiss LM. Toxoplasmosis. Handb Clin Neurol. 2013;114:125–45.

14. Lima TS, Lodoen MB. Mechanisms of Human Innate Immune Evasion by Toxoplasma gondii. Front Cell Infect Microbiol. 2019;9:103.

15. Birgisdóttir A, Asbjörnsdottir H, Cook E, Gislason D, Jansson C, Olafsson I, et al. Seroprevalence of Toxoplasma gondii in Sweden, Estonia and Iceland. Scand J Infect Dis. 2006;38:625–31.

16. Desmettre T. Toxoplasmosis and behavioural changes. J Fr Ophtalmol. 2020;43:e89–93.

17. Saadatnia G, Golkar M. A review on human toxoplasmosis. Scand J Infect Dis. 2012;44:805–14.

18. Dunay IR, Gajurel K, Dhakal R, Liesenfeld O, Montoya JG. Treatment of Toxoplasmosis: Historical Perspective, Animal Models, and Current Clinical Practice. Clin Microbiol Rev. 2018;31:e00057–17.

19. Dupont CD, Christian DA, Hunter CA. Immune response and immunopathology during toxoplasmosis. Semin Immunopathol. 2012;34:793–813.

20. Fabiani S, Pinto B, Bonuccelli U, Bruschi F. Neurobiological studies on the relationship between toxoplasmosis and neuropsychiatric diseases. J Neurol Sci. 2015;351:3–8.

21. Ngoungou EB, Bhalla D, Nzoghe A, Dardé M-L, Preux P-M. Toxoplasmosis and epilepsy--systematic review and meta analysis. PLoS Negl Trop Dis. 2015;9:e0003525.

22. Leroy J, Houzé S, Dardé M-L, Yéra H, Rossi B, Delhaes L, et al. Severe toxoplasmosis imported from tropical Africa in immunocompetent patients: A case series. Travel Med Infect Dis. 2020;35:101509.

23. Simon S, de Thoisy B, Mercier A, Nacher M, Demar M. Virulence of atypical Toxoplasma gondii strains isolated in French Guiana in a murine model. Parasite Paris Fr. 2019;26:60.

24. Shwab EK, Saraf P, Zhu X-Q, Zhou D-H, McFerrin BM, Ajzenberg D, et al. Human impact on the diversity and virulence of the ubiquitous zoonotic parasite Toxoplasma gondii. Proc Natl Acad Sci [Internet]. 2018 [cited 2022 Jun 1];115. Available from: https://pnas.org/doi/full/10.1073/pnas.1722202115

25. Lin R-H, Lai D-H, Zheng L-L, Wu J, Lukeš J, Hide G, et al. Analysis of the mitochondrial maxicircle of Trypanosoma lewisi, a neglected human pathogen. Parasit Vectors. 2015;8:665.

26. Lun Z-R, Wen Y-Z, Uzureau P, Lecordier L, Lai D-H, Lan Y-G, et al. Resistance to normal human serum reveals Trypanosoma lewisi as an underestimated human pathogen. Mol Biochem Parasitol. 2015;199:58–61.

27. Carrera NJR, Carmona MC, Guerrero OM, Castillo AC. [The immunosuppressant effect of T. lewisi (Kinetoplastidae) infection on the multiplication of Toxoplasma gondii (Sarcocystidae) on alveolar and peritoneal macrophages of the white rat]. Rev Biol Trop. 2009;57:13–22.

28. Castillo C, Muñoz L, Carrillo I, Liempi A, Gallardo C, Galanti N, et al. Ex vivo infection of human placental chorionic villi explants with Trypanosoma cruzi and Toxoplasma gondii induces different Toll-like receptor expression and cytokine/chemokine profiles. Am J Reprod Immunol N Y N 1989. 2017;78.

29. Chinchilla M, Guerrero OM, Castro A. Effect of Trypanosoma lewisi infection on the Toxoplasma gondii multiplication in white rat peritoneal macrophages. Parasitol Latinoam. 2004;59:3–7.

30. Chinchilla M, Reyes L, Guerrero O, Castro A. Role of interferon-g on the immunosuppression during Toxoplasma gondii infection by Trypanosoma lewisi. Parasitol Latinoam. 2005;60:54–6.

31. Gao J-M, Yi S-Q, Geng G-Q, Xu Z-S, Hide G, Lun Z-R, et al. Infection with Trypanosoma lewisi or Trypanosoma musculi may promote the spread of Toxoplasma gondii. Parasitology. 2021;148:703–11.

32. Mccown ME, Grzeszak B. Zoonotic and infectious disease surveillance in Central America: Honduran feral cats positive for toxoplasma, trypanosoma, leishmania, rickettsia, and Lyme disease. J Spec Oper Med [Internet]. 2010 [cited 2022 Sep 1];10:41–3. Available from: 10.55460/13SQ-OK4V

33. Dubey JP, Murata FHA, Cerqueira-Cézar CK, Kwok OCH, Su C. Epidemiological Significance of Toxoplasma gondii Infections in Wild Rodents: 2009–2020. J Parasitol [Internet]. 2021 [cited 2022 Jun 1];107. Available from: https://bioone.org/journals/journal-of-parasitology/volume-107/issue-2/20-121/Epidemiological-Significance-of-Toxoplasma-gondii-Infections-in-Wild-Rodents/10.1645/20-121.full

34. Etougbétché JR, Hamidović A, Dossou H-J, Coan-Grosso M, Roques R, Plault N, et al. Molecular prevalence, genetic characterization and patterns of Toxoplasma gondii infection in domestic small mammals from Cotonou, Benin. Parasite Paris Fr. 2022;29:58.

35. Dobigny G, Poirier P, Hima K, Cabaret O, Gauthier P, Tatard C, et al. Molecular survey of rodent-borne Trypanosoma in Niger with special emphasis on T. lewisi imported by invasive black rats. Acta Trop. 2011;117:183–8.

36. Ortiz PA, Garcia HA, Lima L, da Silva FM, Campaner M, Pereira CL, et al. Diagnosis and genetic analysis of the worldwide distributed Rattus-borne Trypanosoma (Herpetosoma) lewisi and its allied species in blood and fleas of rodents. Infect Genet Evol J Mol Epidemiol Evol Genet Infect Dis. 2018;63:380–90.

37. Salzer JS, Pinto CM, Grippi DC, Williams-Newkirk AJ, Peterhans JK, Rwego IB, et al. Impact of Anthropogenic Disturbance on Native and Invasive Trypanosomes of Rodents in Forested Uganda. EcoHealth. 2016;13:698–707.

38. Tatard C, Garba M, Gauthier P, Hima K, Artige E, Dossou DKHJ, et al. Rodent-borne Trypanosoma from cities and villages of Niger and Nigeria: A special role for the invasive genus Rattus? Acta Trop. 2017;171:151–8.

39. Dobigny G, Gauthier P, Houéménou G, Dossou HJ, Badou S, Etougbétché J, et al. Spatiotemporal survey of small mammal-borne Trypanosoma lewisi in Cotonou, Benin, and the potential risk of human infection. Infect Genet Evol. 2019;75:103967.

40. Sikes RS, Gannon WL. Guidelines of the American Society of Mammalogists for the use of wild mammals in research. J Mammal. 2011;92:235–53.

41. Dossou H-J, Le Guyader M, Gauthier P, Badou S, Etougbetche J, Houemenou G, et al. Finescale prevalence and genetic diversity of urban small mammal-borne pathogenic Leptospira in Africa: A spatiotemporal survey within Cotonou, Benin. Zoonoses Public Health. 2022;

42. Badou S, Hima K, Agbangla C, Gauthier P, Missihoun AA, Houéménou G, et al. Biological invasions in international seaports: a case study of exotic rodents in Cotonou. Urban Ecosyst [Internet]. 2023 [cited 2023 Jun 24]; Available from: 10.1007/s11252-023-01356-6

43. Dossou H-J, Tenté B, Houémènou G, Sossou MD, Rossi J-P, Dobigny G. Fine-scale Landscape Variability of Cotonou City, Benin: Insights From Three Contrasted Urban Neighborhoods. 2021;

44. Ségard A, Romero A, Ravel S, Truc P, Dobigny G, Gauthier P, et al. Development of nine microsatellite loci for Trypanosoma lewisi, a potential human pathogen in Western Africa and South-East Asia, and preliminary population genetics analyses. Peer Community J [Internet]. 2022 [cited 2022 Dec 15];2:e69. Available from: https://peercommunityjournal.org/articles/10.24072/pcjournal.188/

45. Gotelli NJ. Null model analysis of species co-occurrence patterns. Ecology. 2000;81:2606–21.

46. Gotelli NJ, Entsminger GL. Swap algorithms in null model analysis. Ecology. 2003;84:532–5.

47. Ulrich W, Gotelli NJ. Null model analysis of species associations using abundance data. Ecology. 2010;91:3384–97.

48. Ulrich W. Pairs—a FORTRAN program for studying pair-wise species associations in ecological matrices. URL Www Keib Umk Plpairs. 2008;

49. Stone L, Roberts A. The checkerboard score and species distributions. Oecologia. 1990;85:74–9.

50. Anderson DR, Burnham KP. Avoiding pitfalls when using information-theoretic methods. J Wildl Manag. 2002;912–8.

51. Team RC. A language and environment for statistical computing. 2015. R foundation for statistical computation, Vienna, Austria. 2020.

52. Bates D, Kliegl R, Vasishth S, Baayen H. Parsimonious mixed models. ArXiv Prepr ArXiv150604967. 2015;

53. Barton K, Barton MK. Package ‘mumin.’ Version. 2015;1:439.

54. Cassan C, Diagne CA, Tatard C, Gauthier P, Dalecky A, Bâ K, et al. Leishmania major and Trypanosoma lewisi infection in invasive and native rodents in Senegal. PLoS Negl Trop Dis. 2018;12:e0006615.

55. Archer CE, Schoeman MC, Appleton CC, Mukaratirwa S, Hope KJ, Matthews GB. Predictors of Trypanosoma lewisi in Rattus norvegicus from Durban, South Africa. J Parasitol. 2018;104:187–95.

56. Garcia HA, Rangel CJ, OrtÍz PA, Calzadilla CO, Coronado RA, Silva AJ, et al. Zoonotic Trypanosomes in Rats and Fleas of Venezuelan Slums. EcoHealth. 2019;16:523–33.

57. Dubey JP, Ferreira LR, Alsaad M, Verma SK, Alves DA, Holland GN, et al. Experimental Toxoplasmosis in Rats Induced Orally with Eleven Strains of Toxoplasma gondii of Seven Genotypes: Tissue Tropism, Tissue Cyst Size, Neural Lesions, Tissue Cyst Rupture without Reactivation, and Ocular Lesions. Knoll LJ,editor. PLOS ONE [Internet]. 2016 [cited 2022 Jun 1];11:e0156255. Available from: https://dx.plos.org/10.1371/journal.pone.0156255

58. Houéménou, Gauthier, Etougbeé tché, Badou, Dossou Agossou, et al. Pathogenic Leptospira in Commensal Small Mammals from the Extensively Urbanized Coastal Benin. Urban Sci [Internet]. 2019 [cited 2022 Dec 15];3:99. Available from: https://www.mdpi.com/2413-8851/3/3/99

59. Ahmed N, French T, Rausch S, Kühl A, Hemminger K, Dunay IR, et al. Toxoplasma coinfection prevents Th2 differentiation and leads to a helminth-specific Th1 response. Front Cell Infect Microbiol. 2017;7:341.

60. Miller CMD, Smith NC, Ikin RJ, Boulter NR, Dalton JP, Donnelly S. Immunological interactions between 2 common pathogens, Th1-inducing protozoan Toxoplasma gondii and the Th2-inducing helminth Fasciola hepatica. PloS One. 2009;4:e5692.

61. Neal LM, Knoll LJ. Toxoplasma gondii profilin promotes recruitment of Ly6Chi CCR2+ inflammatory monocytes that can confer resistance to bacterial infection. PLoS Pathog. 2014;10:e1004203.

62. Souza MC, Fonseca DM, Kanashiro A, Benevides L, Medina TS, Dias MS, et al. Chronic Toxoplasma gondii Infection Exacerbates Secondary Polymicrobial Sepsis. Front Cell Infect Microbiol. 2017;7:116.

63. Thomasson D, Wright EA, Hughes JM, Dodd NS, Cox AP, Boyce K, et al. Prevalence and coinfection of Toxoplasma gondii and Neospora caninum in Apodemus sylvaticus in an area relatively free of cats. Parasitology. 2011;138:1117–23.

64. Watanabe H, Suzuki Y, Makino M, Fujiwara M. Toxoplasma gondii: induction of toxoplasmic encephalitis in mice with chronic infection by inoculation of a murine leukemia virus inducing immunodeficiency. Exp Parasitol. 1993;76:39–45.

